# Pegivirus avoids immune recognition but does not attenuate acute-phase disease in a macaque model of HIV infection

**DOI:** 10.1101/175646

**Authors:** Adam L. Bailey, Connor R. Buechler, Daniel R. Matson, Eric J. Peterson, Kevin G. Brunner, Mariel S. Mohns, Meghan Breitbach, Laurel M. Stewart, Adam J. Ericsen, Saverio Capuano, Heather A. Simmons, David T. Yang, David H. O’Connor

**Author notes:** These authors contributed equally to this work (Adam L. Bailey, Connor R. Buechler). To whom correspondence should be addressed: Dr. Adam Bailey.

## Abstract

Human pegivirus (HPgV) protects HIV+ people from HIV-associated disease, but the mechanism of this protective effect remains poorly understood. We sequentially infected cynomolgus macaques with simian pegivirus (SPgV) and simian immunodeficiency virus (SIV) to model HIV+HPgV co-infection. SPgV had no effect on acute-phase SIV pathogenesis – as measured by SIV viral load, CD4+ T cell destruction, and immune activation – suggesting that HPgV’s protective effect is exerted primarily during the chronic phase of HIV infection. We also examined the immune response to SPgV in unprecedented detail, and found that this virus elicits virtually no activation of the immune system despite persistently high titers in the blood over long periods of time. Overall, this study expands our understanding of the pegiviruses – an understudied group of viruses with a high prevalence in the global human population – and suggests that the protective effect observed in HIV+HPgV co-infected people occurs primarily during the chronic phase of HIV infection.

**One Sentence Summary:** Pegivirus avoids immune recognition but does not attenuate acute-phase disease in a macaque model of HIV infection.

**Short Title:** Pegivirus and AIDS-virus co-infection

**Accessible Summary:** People infected with HIV live longer, healthier lives when they are co-infected with the human pegivirus (HPgV) – an understudied virus with a high prevalence in the global human population. To better understand how HPgV protects people with HIV from HIV-associated disease, we infected macaques with simian versions of these two viruses (SPgV and SIV). We found that SPgV had no impact on the incidence of SIV-associated disease early during the course of SIV infection – a time when SIV and HIV are known to cause irreversible damage to the immune system. Oddly, we found that the immune system did not recognize SPgV; a finding that warrants further investigation. Overall, this study greatly expands on our understanding of the pegiviruses and their interaction with the immune system.

## Introduction

Human pegivirus (HPgV) – formerly known as GB virus C (GBV-C) and also as Hepatitis G Virus (HGV) – is a positive-sense, single-stranded RNA virus in the Pegivirus genus of the *Flaviviridae* family (*1*). HPgV infects one out of six humans globally and is frequently transmitted via blood products (*2*). Little is known about the molecular biology of pegiviruses and the natural course of HPgV infection is poorly understood. However, HPgV causes persistent, high-titer viremia without eliciting symptoms or overt signs of disease (*3*, *4*). Interestingly, epidemiological studies have found that people infected with human immunodeficiency virus (HIV) experience reduced disease when they are co-infected with HPgV. Specifically, HIV-infected individuals co-infected with HPgV are protected from HIV-induced CD4 T cell depletion (*5-8*) and pathological immune activation (*9-12*). These individuals also experience a 2.5-fold reduction in all-cause mortality relative to HIV+ individuals not co-infected with HPgV (see (*13*) for a meta-analysis and (*2*) for a review). However, the timing and mechanistic underpinnings of this protective association are not known, in part because most data on HIV+HPgV co-infection comes from cross-sectional studies performed during the chronic phase of HIV infection. In particular, the impact of HPgV infection on acute phase HIV infection – a period during which pathologic changes in the HIV-infected host are most dramatic (*14*) – has not been studied. As such, the impact of HPgV co-infection on the natural course of HIV infection, and the mechanism(s) by which HPgV attenuates HIV disease *in vivo,* remain uncharacterized.

Macaque monkeys infected with simian immunodeficiency virus (SIV) exhibit several features of progressive HIV disease in humans, including CD4+ T cell depletion and pathological immune activation. As such, macaques infected with SIV are a valuable model for investigating the pathogenesis of HIV *in vivo* (*15*). We recently discovered simian pegiviruses (SPgVs) infecting wild baboons in Africa (*16*) and used blood from an olive baboon (*Papio anubis*) sampled in Kibale, Uganda to infect captive cynomolgus macaques (*Macaca fascicularis*) with SPgV. This resulted in the first laboratory-animal model of HPgV infection (*17*). Notably, SPgV infection causes persistent, high-titer viremia in macaques without eliciting signs of disease, recapitulating several defining features of HPgV infection in humans.

Here, we used SPgV and SIV infection in Mauritian cynomolgus macaques to model HPgV and HIV co-infection in humans. We compared SIV disease parameters in four SPgV+SIV co-infected macaques to four macaques infected with SIV-only, with the hypothesis that SPgV would attenuate SIV pathogenesis during the acute phase of SIV infection, or result in improved recovery from acute insult of SIV infection, as is observed in natural simian hosts of SIV which are often co-infected with their own species-specific SPgVs (*18*, *19*).

## Results

### SPgV pre-infection does not impact acute-phase SIV viremia

For this study, eight female cynomolgus macaques (*Macaca fascicularis*) with identical major histocompatibility complex (MHC) class I haplotypes (M3/M4) were randomized to receive SPgV or no intervention prior to SIV infection (Fig S1, Table S1). Four macaques received intravenous inoculations of plasma containing 2.29×10^7^ genome copies (gc) of SPgV measured by quantitative RT-PCR, as done previously (*17*). These infections achieved initial peak titers between 7.30×10^6^ and 2.66×10^7^ gc/ml of plasma between 7 and 14 days post SPgV infection (Fig. 1A), similar to what had been observed previously. By 26 days post-SPgV infection viral loads had established a high-titer steady-state (average of 2.02×10^7^ gc/ml) and all eight macaques were inoculated intra-rectally with 7,000 tissue-culture-infectious-dose (TCID)_50_ of SIVmac239.

**Figure 1.**
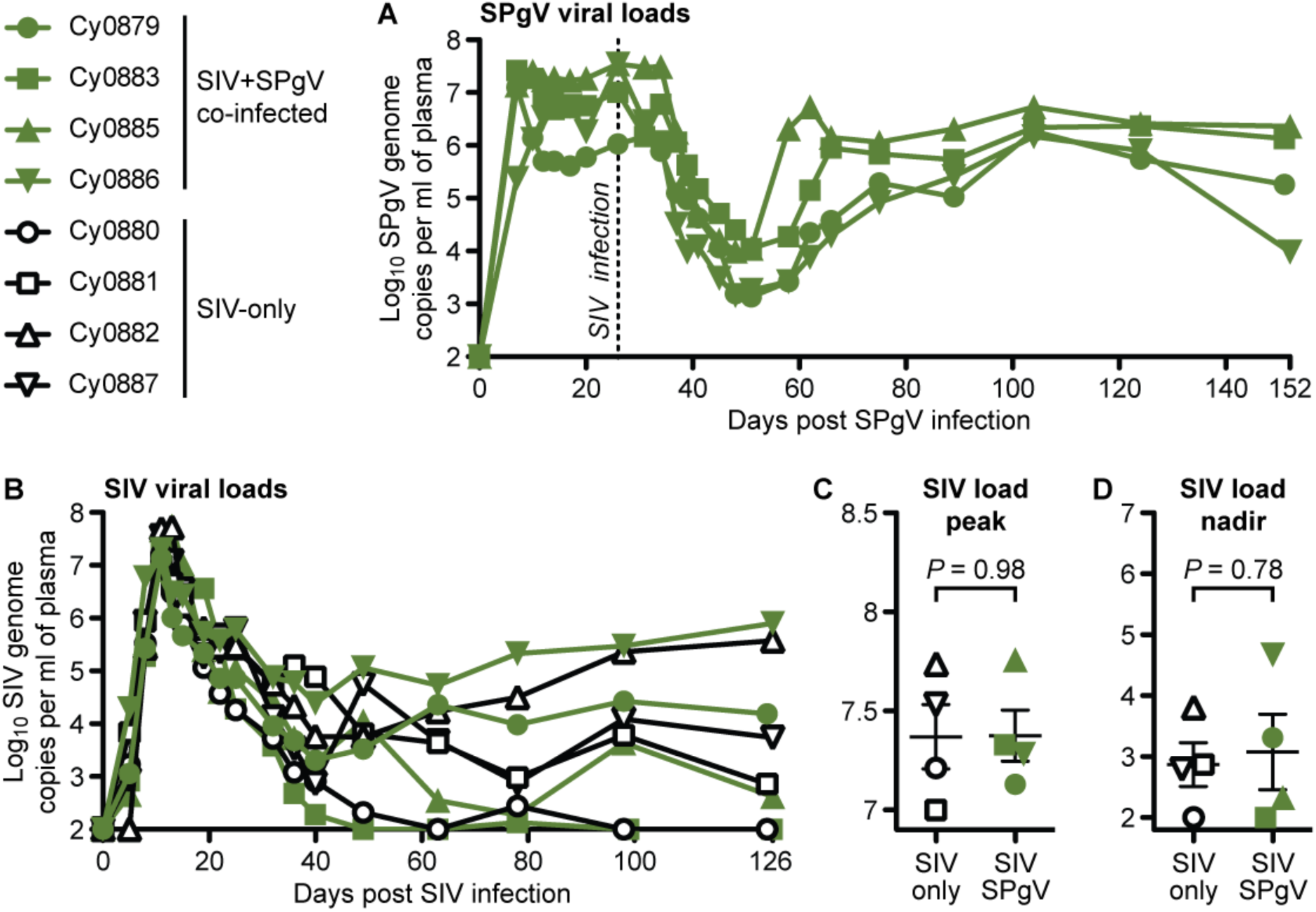
SPgV and SIV viral loads in infected macaques. Titers for each virus were measured from plasma using highly sensitive virus-specific quantitative RT-PCR assays. (**A**) SPgV titers in the four macaques infected with SPgV+SIV. (**B**,**C**,**D**) SIV titers in four macaques infected with SPgV+SIV (green) and four macaques infected with SIV-only (black). *P* values reflect a two-tailed unpaired t-test and error bars represent SEM. The symbols used for each animal in this figure are consistent throughout the manuscript.

SIV plasma viral loads followed a typical trajectory during acute phase, reaching peak titers between 1.01×10^7^ and 5.25×10^7^ gc/ml of plasma between days 11 and 13 post-SIV infection in all eight macaques (Fig. 1B). No differences in peak SIV plasma titer or post-peak nadir were observed between the SIV+SPgV co-infected and SIV-only groups (Fig. 1C). To determine whether SPgV impacted subsequent SIV viral load trajectory, we followed the macaques for 126 days after SIV infection. Within each group, we observed a wide range of viral load set-point titers. However, there was not a significant difference in SIV viral loads between the SIV+SPgV group and the SIV-only group at any time point (Fig 1D).

### Acute SIV infection reduces SPgV viremia

Beginning as early as day 5 of SIV infection (day 31 of SPgV infection), we observed a significant drop in SPgV plasma viral loads in all SPgV+SIV co-infected macaques. The decline in SPgV viral loads reached a nadir of 1.36×10^3^ - 1.11×10^4^ gc/ml of plasma between day 22 and 25 of SIV infection (day 48-51 of SPgV infection), then gradually rebounded to a new set-point that was approximately 1.5 log_10_ lower than the pre-SIV viremic set-point by day 40-49 of SIV infection (66-75 of SPgV infection). Previously, we showed that SPgV accumulates little-to-no sequence variation over time in infected macaques. Therefore, we deep sequenced the SPgV genome from each animal before the decline (day -6 of SIV infection; day 20 SPgV infection) and after recovery of high-titer SPgV viremia (day 49 of SIV infection; day 75 SPgV infection) to look for unique signatures of immune pressure on SPgV that may have been triggered by SIV infection. While SPgV from two macaques accumulated 1-3 protein-coding (*i.e.* non-synonymous) mutations over this period, SPgV from the other two macaques revealed no protein-coding mutations (Table S2), suggesting that an SPgV-specific immune response and subsequent mutational escape was not responsible for the measured decrease in SPgV viral loads.

We hypothesized that the reduction in SPgV viral load during acute SIV infection was the result of inflammation induced by SIV, which has been reported to occur secondary to microbial translocation from the gut lumen (*14*). Thus, we infected eight macaques intravenously with 2.29×10^7^ gc of SPgV and treated four of these macaques with dextran sulfate sodium (DSS) on day 26 post-SPgV infection to induce microbial translocation (*20*, *21*). DSS treatment had no significant impact on SPgV viral loads, suggesting that inflammation caused by microbial translocation during acute SIV infection was not responsible for the observed decline in SPgV viral loads (Table S3).

### SPgV pre-infection does not prevent loss of peripheral CD4+ T cells

The absolute number of circulating CD4+ T cells remains the most clinically-relevant marker of HIV/SIV disease progression, and some studies of HIV+ human patients have shown a modest association between HPgV co-infection and higher peripheral CD4+ T cell counts (*5-8*). Therefore, we followed blood CD4+ T cell counts in both the SIV-only and SIV+SPgV groups. We observed an initial drop in the circulating CD4+ T cells that corresponded with peak SIV viremia in all eight animals which then recovered to near pre-SIV levels. However, there were no statistically significant differences in the CD4+ T cell count between the SIV-only and SIV+SPgV groups (Fig. 2A). An increase in the absolute number of circulating CD4+ T cells was observed prior to SIV infection in the macaques infected with SPgV, although a concomitant increase was also noted in the SPgV-naïve macaques during this time period, suggesting that this increase was not due to SPgV infection.

**Figure 2.**
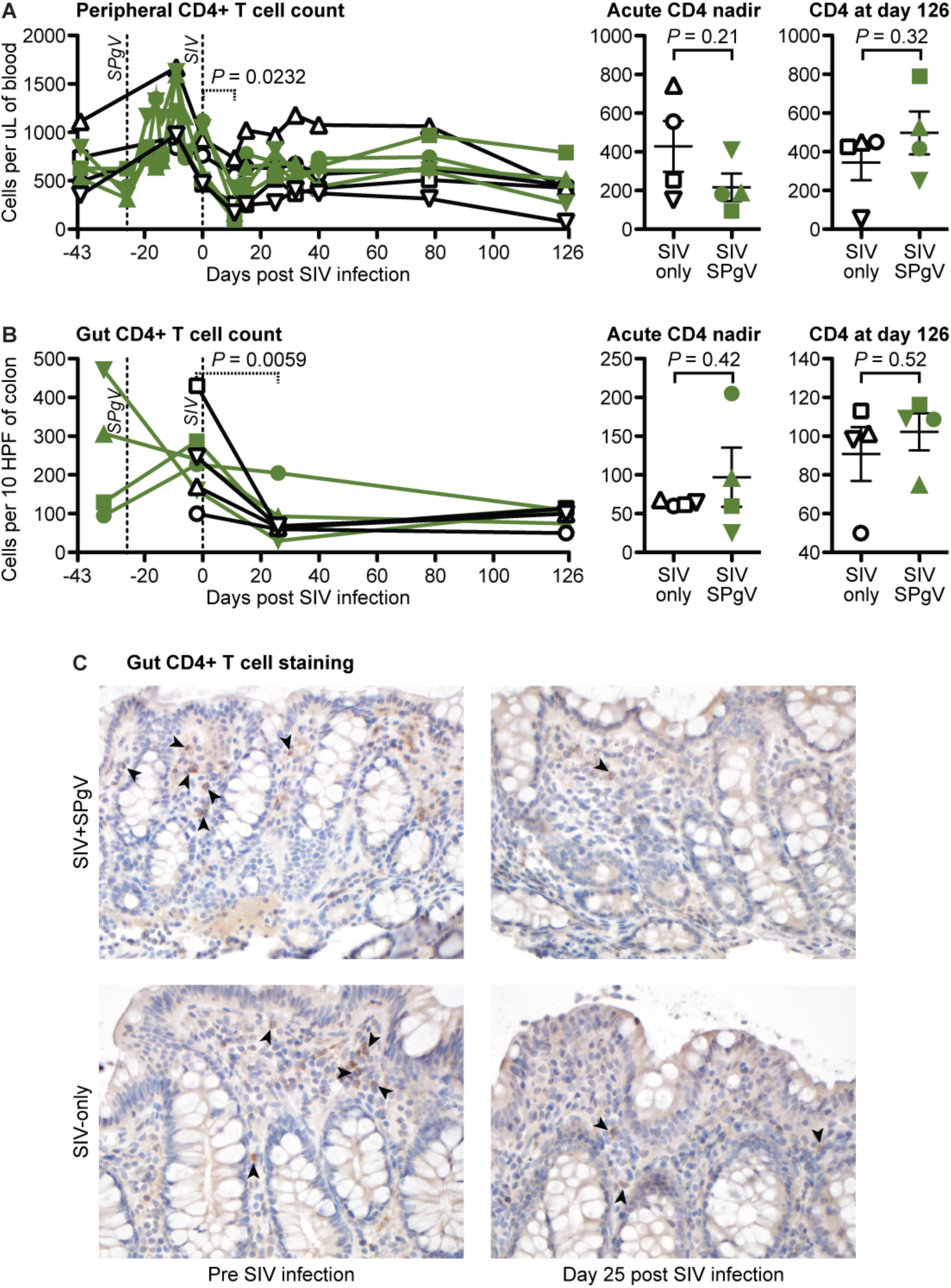
SIV pathogenesis in SIV-only vs. SIV+SPgV infected macaques. (**A**) Peripheral CD4+ T cell counts were obtained by multiplying absolute lymphocyte counts by the percentage of lymphocytes that were CD3+ CD4+ CD20-CD8- (see Figure 3 for gating strategy details). (**B**) Gut CD4+ T cells were stained within sections of colonic tissues via IHC with an anti-CD4 antibody and manually quantified. Significant differences between the SIV-only and SPgV+SIV groups were analyzed using a two-tailed unpaired t-test (solid line) with error bars representing SEM. Significant changes in all animals over the course of acute SIV infection were quantified using a two-tailed paired t-test (dashed line). (**C**) A representative set of colonic tissue from Cy0883 (SIV+SPgV) and Cy0887 (SIV-only) are shown pre and post SIV infection at 400x for comparison. Arrows highlight representative cells with membranous CD4 staining.

### SPgV pre-infection does not prevent loss of gut CD4+ T cells

The early loss of CD4+ T cells in the gastrointestinal tract is a hallmark of HIV/SIV pathogenesis (*14*, *22*); yet the effect of HPgV/SPgV co-infection on gut CD4+ T cell depletion has never been examined. To see if SPgV pre-infection protected gut CD4+ T cells from SIV-mediated destruction, we collected colon pinch-biopsies from animals pre-and post-SIV infection, then analyzed the abundance of lamina propria CD4+ cells using immunohistochemistry (IHC). As expected, SIV infection led to an acute loss of gut-resident CD4+ cells, but the loss in the SIV+SPgV group was not statistically different compared to the SIV-only group (Fig. 2B,C).

**Figure 3.**
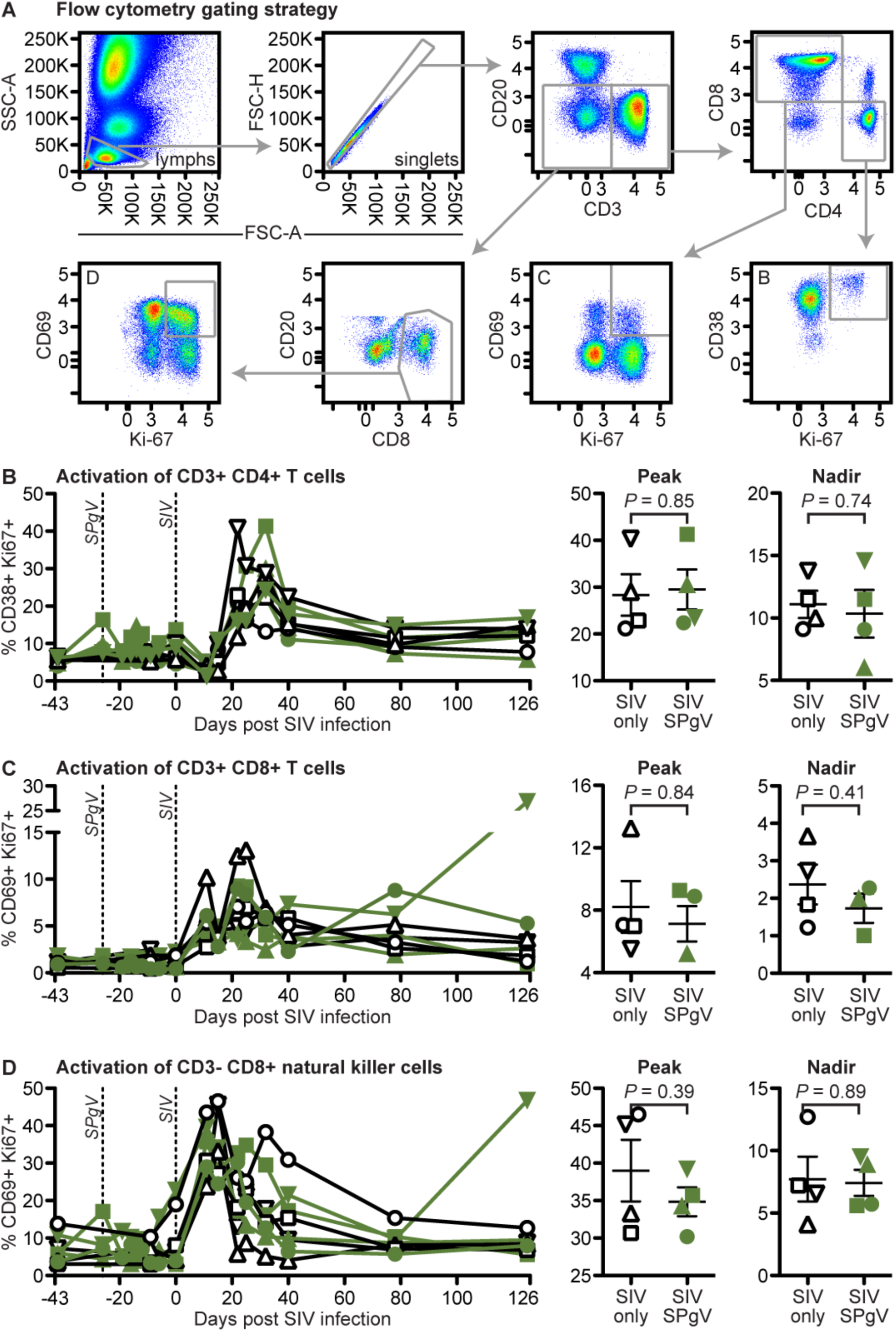
Peripheral immune activation in SIV-only vs. SIV+SPgV infected macaques. Fresh whole blood was used for staining and flow cytometry at each timepoint. *P* values represent a two-tailed unpaired t-test with error bars reflecting SEM. Note: Cy0886 did not exhibit a distinct peak or nadir of CD69+ Ki67+ expression in the CD3+ CD8+ T cell population, and so is not included in these analyses.

### SPgV pre-infection does not reduce pathological immune activation during acute SIV infection

Several studies have demonstrated a correlation between HPgV infection and reduced immune activation in HIV-infected people (*9-12*). However, none of these studies have examined the effect of HPgV pre-infection on the trajectory of HIV disease during acute HIV infection, a time when HIV is known to cause irreversible damage to the immune system. Therefore, we trended changes in peripheral immune cell activation after SIV infection using flow cytometry. For each immune cell subset examined (CD3+CD4+ T cells, CD3+CD8+ T cells, and CD3-CD8+ natural killer [NK] cells) we chose a combination of activation markers that most clearly delineated a positive and negative population (Fig 3A). While the timing and magnitude of activation following SIV infection varied by cell subset, we did not detect a significant difference in the magnitude of peak activation, the time to peak activation, or the post-peak nadir of activation between the SIV-only and SIV+SPgV groups within any subset (Fig. 3B-D). Activation of immune cells in the gut and in lymph nodes, as measured by IHC staining for the proliferation marker Ki67, showed a similar pattern (Fig. 4).

**Figure 4.**
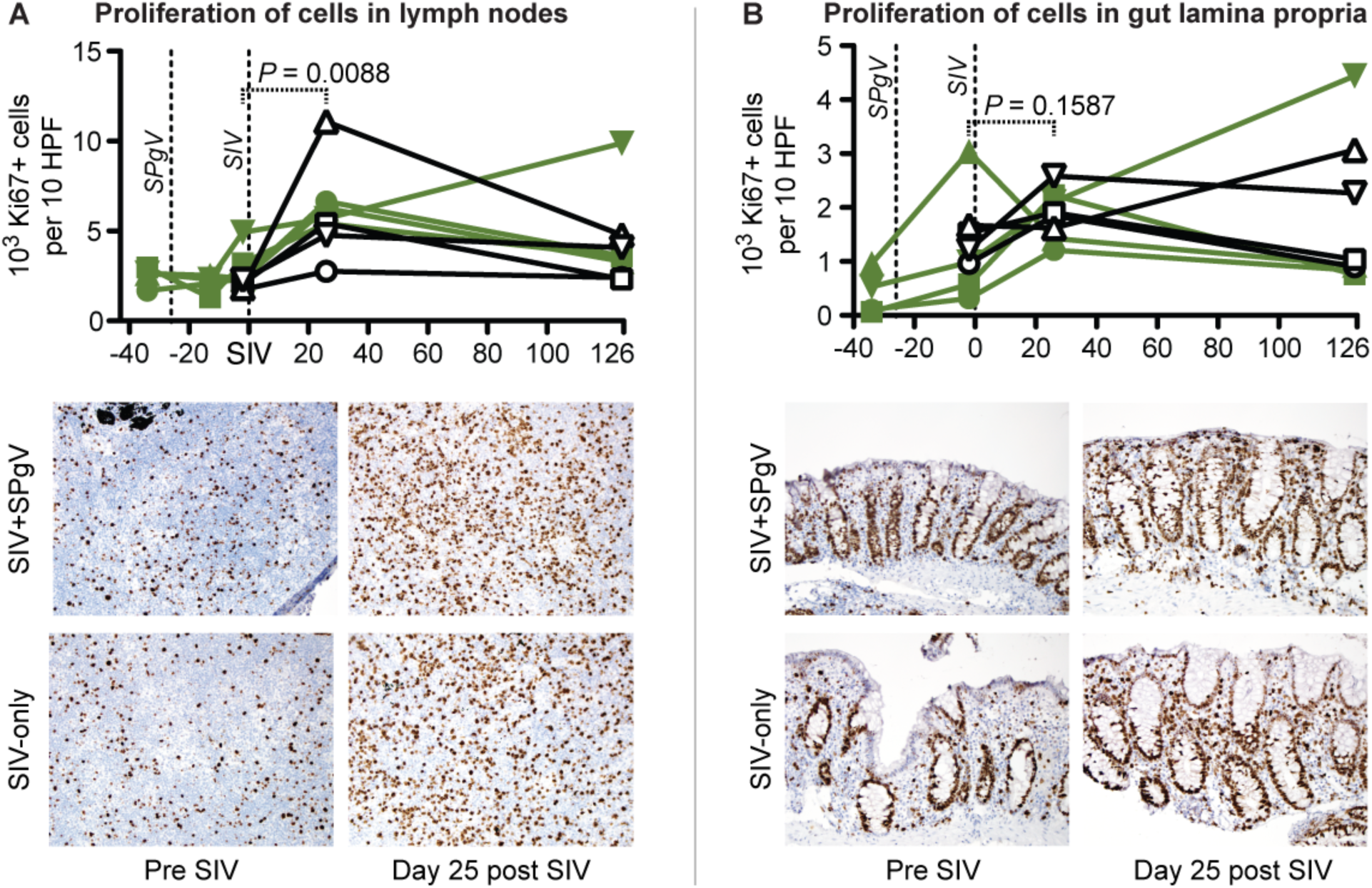
Activation of immune tissues in SIV-only vs. SIV+SPgV infected macaques. Proliferating cells were stained within sections of lymph nodes (**A**) and colon (**B**) via IHC with an anti-Ki67 antibody and manually quantified. Significant changes over time were quantified using a two-tailed paired t-test (dashed line). A representative set of lymph nodes from Cy0883 (SIV+SPgV) and Cy0881 (SIV-only) is shown at 400X pre and post SIV infection for comparison in (**A**). A representative set of colon tissues from Cy0886 (SIV+SPgV) and Cy0887 (SIV-only) is shown pre and post SIV infection at 400x for comparison in (**B**).

### SPgV infection does not induce activation of the immune system

Systemic viral infections typically elicit a Th1-type immune response characterized by the activation of lymphocytes. To examine the immune response to acute pegivirus infection, we trended markers of immune cell activation by flow cytometry for 26 days following SPgV infection. Oddly, we saw no significant changes from pre-infection baseline in the total number or activation state of circulating lymphocytes during this time period despite high titers of SPgV. This is in stark contrast to SIV infection, which elicited a robust increase in immune activation during the same timeframe (Fig. 5).

**Figure 5.**
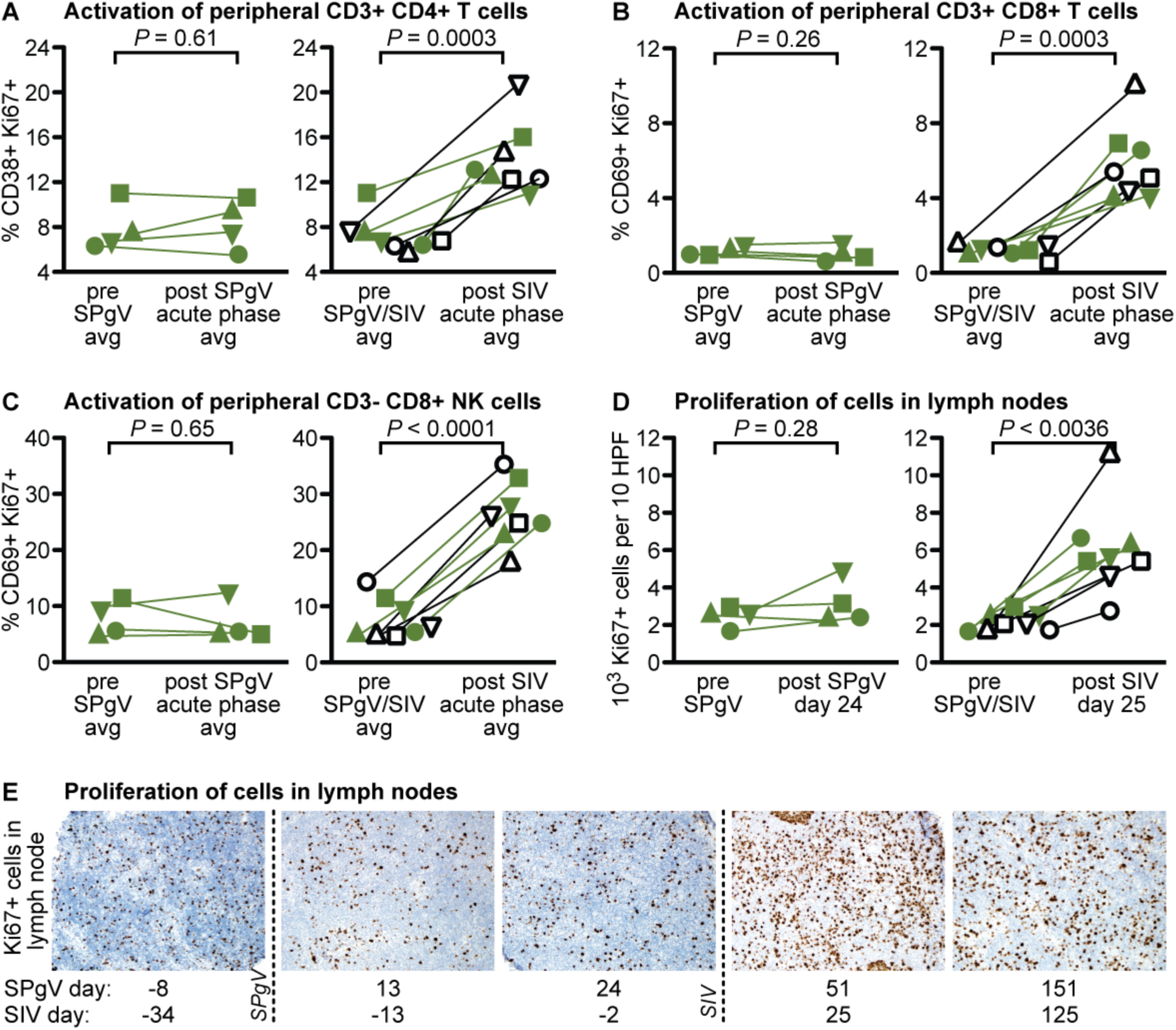
Immune activation following SPgV vs. SIV infection. (**A-C**) Fresh whole blood was used at each timepoint. *P* values are from a two-tailed paired t-test comparing pre-infection average to post-SPgV or post-SIV average within the first 26 days post-infection for each virus, respectively. (**D**) Proliferating cells were stained within sections of lymph nodes via IHC with an anti-Ki67 antibody and manually quantified. Significant changes in over time were quantified using a two-tailed paired t-test. (**E**) Representative set of lymph node tissue from Cy0885 is shown at 400x.

## Discussion

HIV+ people experience slower disease progression and reduced mortality when co-infected with HPgV, but the mechanisms by which HPgV mediates this protective effect are not known. Here, we utilized a recently-developed macaque model to study SPgV+SIV co-infection and found that pre-infection with SPgV had no effect on acute-phase SIV pathogenesis as measured by SIV viral loads, peripheral and gut-resident CD4+ T cell depletion, or SIV-induced immune activation. One interpretation of these findings could be that, unlike HIV+HPgV co-infection in humans, SPgV does not protect macaques from SIV disease. While this is possible, studies of SPgVs in non-human primates have shown that the biology of these viruses closely mirrors HPgV in humans (*1*, *16*, *17*, *19*, *23*). The progression of SIV infection in macaques also closely mirrors that of HIV infection in humans, and so it appears that SIV+SPgV infection in macaques is a close approximation of HIV+HPgV in humans. Therefore, our data suggest that HPgV does not exert a protective effect on HIV pathogenesis during the acute phase of HIV infection, but rather alters HIV/AIDS disease during the chronic phase. This conclusion is supported by some of the epidemiologic data on HIV+HPgV co-infection (*8*, *13*). Unfortunately, confirming and studying an effect on this timescale (*i.e.* years) is intractable in macaques. Nonetheless, a protective effect in the chronic phase of HIV infection may ultimately prove to be of greater clinical value, should a HPgV-inspired anti-HIV therapy ever come to fruition.

Our study resulted in the first-ever, highly-detailed analysis of the immune response to acute pegivirus infection. Previous studies of persistent pegivirus infection showed that these viruses accumulate very few mutations over time, which is in stark contrast to other persistent RNA viruses like HIV/SIV, hepatitis C virus, and simian arteriviruses (*17*, *24-26*). These viruses rapidly acquire mutations that alter protein structure, allowing them to escape adaptive host immune responses targeting viral proteins (*27-29*). Thus, the lack of sequence changes observed in the pegivirus genome over the course of infection alludes to an uncharacteristically weak or absent host immune response. To investigate this further, we measured the activation of immune cells following SPgV infection. While SIV infection resulted in robust activation of all cell subsets examined, we found that SPgV infection elicited no detectable activation of the immune system. This is the first direct *in vivo* evidence demonstrating the absence of an anti-pegivirus immune response. It remains to be determined whether pegiviruses inhibit activation of the immune system or simply avoid immune detection, but answering this question will require the development of additional virus-specific reagents and assays. Interestingly, either scenario would require that these viruses employ a unique mechanism to maintain high-titer, persistent infection in the primate host.

Although pegivirus infection does not appear to elicit a substantial immune response, the significant drop in SPgV plasma viral loads that we observed upon co-infection of macaques with SIV suggests that pegivirus replication may be restricted by the immune activation which occurs as a result of acute SIV infection. One of the major drivers of immune activation during early SIV/HIV infection is the translocation of microbial products from the gut lumen into systemic circulation, which occurs as a result of the destruction of gut-resident CD4+ T cells. To determine whether this impacted SPgV replication, we treated SPgV infected macaques with DSS, a chemical known to induce microbial translocation. These macaques experienced no decrease in SPgV plasma viral loads, in contrast to SPgV+ monkeys infected with SIV. This possibly suggests that antiviral innate immune factors induced by SIV were responsible for decreased SPgV replication during the acute-phase of SIV infection. Alternatively, SPgV and SIV could share the same target cell-type, and the destruction of these cells by SIV could explain the temporary reduction in SPgV plasma viral loads. This hypothesis warrants further investigation, and studies designed to determine the tissue tropism of the pegiviruses and SIV/HIV are ongoing. We are hopeful that future pegivirus studies will provide a deeper understanding of the biology of these enigmatic viruses, and ultimately the mechanisms by which HPgV protects humans from HIV-associated disease.

## Materials and Methods

### Care and use of animals

All macaque monkeys used in this study were cared for by the staff at the Wisconsin National Primate Research Center in accordance with the regulations and guidelines outlined in the Animal Welfare Act and the Guide for the Care and Use of Laboratory Animals. Details of this study (UW-Madison Animal Care and Use Protocol No. G00707) were approved by the University of Wisconsin Institutional Animal Care and Use Committee, in accordance with recommendations of the Weatherall Report.

### Selection of animals

To control for host genetic factors to the extent possible, we used cynomolgus macaques from the island of Mauritius, where there is an inbred macaque population due to a recent founder effect. All animals selected for this study were female and were major histocompatibility complex (MHC)-matched: all animals were heterozygous with an M3/M4 combination of MHC haplotypes. Unlike other defined MHC haplotypes found in Mauritian cynomolgus macaques (*e.g.* M1), spontaneous control of SIV infection is not known to be associated with the M3 or M4 haplotype (*30*).

### Virus inoculations

A SPgV stock was created for this study by aliquoting plasma collected from a macaque (cy0500) inoculated intravenously with plasma from an SPgV+ olive baboon (*Papio anubis*) sampled in Kibale National Park, Uganda (GenBank accession: KF234530), as described in detail previously (*16*, *17*). Macaques infected with SPgV in this study were inoculated intravenously with 700 μL of cy0500 plasma containing 2.29 × 10^7^ genome copies of SPgV. SIV infections were achieved using a single intrarectal inoculation of 7,000 tissue-culture-infectious-dose (TCID)_50_ of the molecularly cloned SIVmac239 virus (GenBank accession: M33262).

### Chemical induction of microbial translocation

Microbial translocation was induced as described previously (*21*). Briefly, a 0.5% solution of dextran sulfate sodium (DSS) was prepared by resuspending colitis-grade DSS (MPBio, Santa Ana, CA) in sterile drinking water and stored at 4°C. Animals were treated once per day for 5-consecutive days with 200 mL of the DSS-containing drinking water, administered by gavage. Animals were monitored for clinical signs of colitis and gastrointestinal distress, and received palliative and clinical care at the full discretion of WNPRC veterinarians.

### RNA extraction from plasma for sequencing and viral loads

RNA was extracted from 300 μL of EDTA-treated plasma using the Viral Total Nucleic Acid Purification Kit (Promega, Madison, WI) on a Maxwell 16 MDx instrument and eluted in 50 μL of DNAse/RNAse free water.

### SPgV Viral loads

A Taqman quantitative RT-PCR (qRT-PCR) assay was used to quantify viral RNA for SPgV (forward primer: 5’-CGGTGTTCATGGCAGGTAT-3’; reverse primer: 5’-CAGTTACAGCCGCGTGTTT-3’; probe: 5’-6FAM-ATGCACCCTGATGTAAGCTGGGCAA-BHQ1-3’), as described previously (*16*). Reverse transcription and PCR were performed using the SuperScript II One-Step qRT-PCR system (Invitrogen, Carlsbad, CA) on a LightCycler 480 instrument (Roche, Indianapolis, IN). Reverse transcription was carried out at 37°C for 15 minutes and then 50°C for 30 minutes followed by 2 minutes at 95°C, and then 50 cycles of amplification as follows: 95°C for 15 sec and 60°C for 1 minute. The 20 μL reaction mixture contained 5 μL of extracted RNA, MgSO_4_ at a final concentration of 3.0 mM, with the two amplification primers at a concentration of 500 nM and probe at a concentration of 100 nM. RNA copy number was calculated using a standard curve that was sensitive down to 10 copies of RNA transcript per reaction.

### SIV viral loads

A Taqman qRT-PCR assay was used to quantify viral RNA for SIV (forward primer: 5’-GTCTGCGTCATCTGGTGCATTC-3’; reverse primer: 5’-CACTAGCTGTCTCTGCACTATGTGTTTTG-3’; probe: 5’-6FAM-CTTCCTCAGTGTGTTTCACTTTCTCTTCTGCG-BHQ1-3’), as described previously (*31*). Cycling conditions were: 37°C for 15 min, 50°C for 30 min, and 95°C for 2 min, followed by 50 amplification cycles of 95°C for 15 sec and 62°C for 1 min with ramp times set to 3°C/sec. The final reaction mixtures (20 μL total volume) contained 0.2 mM dNTPs, 3.5 mM MgSO4, 150 ng random hexamer primers (Promega, Madison, WI), 0.8 μL SuperScript II One-Step qRT-PCR enzyme mix, 600 nM of each amplification primer and 100 nM of probe.

### Amplicon-based sequencing of SPgV

SPgV was amplified with the Qiagen OneStep RT-PCR kit and five overlapping ∼2.5kb amplicons. Primers were designed using Primer3 (*32*) in Geneious R9 (Biomatters, Auckland, NZ) (Table S4). Cycling conditions were: 50°C for 30 minutes and 94°C for 2 minutes, followed by 40 cycles of 94°C for 15 seconds, 55°C for 30 seconds, and 68°C for 2.5 minutes, followed by a final extension at 68°C for 5 minutes. Amplicons were fragmented and sequencing adaptors were added using the Nextera DNA Sample Preparation Kit (Illumina, San Diego, CA). Deep sequencing was performed on the Illumina MiSeq, and sequence data were analyzed using Geneious Pro R9 (Biomatters, Auckland, NZ). Low quality (<Q40, Phred quality score) and short reads [<100 base pairs (bp)] were removed, and coding-complete genome sequences for SPgV were acquired by mapping reads to the reference sequence for the SPgVkob inoculum (Genbank ID KF234530) using the Geneious alignment tool at medium-low sensitivity. Consensus SPgV sequences from each animal at each timepoint were compared to the inoculum using a ClustalW alignment with an IUB cost matrix, gap open cost of 15, and gap extend cost of 6.66.

### Immune cell activation and CD4+ T cell counts

Staining for flow cytometry was performed on EDTA-anticoagulated whole blood as described previously (Pomplun, 2015). Briefly, 0.1 mL of EDTA-anticoagulated whole-blood was incubated for 15 min at room temperature in the presence of a mastermix of antibodies against CD38 (clone AT1, FITC conjugate, 20 μl), CD69 (clone TP1.55.3, ECD conjugate, 10 μL), CD3 (clone SP34-2, Alexa Fluor 700 conjugate, 3 μl), CD25 (clone M-A251, Brilliant Violet 421 conjugate, 5 μl), CD8 (clone SK1, Brilliant Violet 510 conjugate, 2.5 μl), CD20 (clone 2H7, Brilliant Violet 650 conjugate, 2 μl), CD4 (clone L200, Brilliant Violet 711 conjugate, 5 μl) antigens. All antibodies were obtained from BD BioSciences (San Jose, CA, USA), except the CD69-specific antibody, which was purchased from Beckman Coulter (Brea, CA, USA) and the CD38-specific antibody, which was purchased from Stemcell Technologies (Vancouver, BC, Canada). Cells were also stained with LIVE/DEAD Fixable Near-IR during this time (Invitrogen, Carlsbad, CA). Red blood cells were lysed using BD Pharm Lyse, after which they were washed twice in media and fixed with 0.125 mL of 2% paraformaldehyde for 20 min. After an additional wash the cells were permeabilized using Bulk Permeabilization Reagent from Invitrogen (Carlsbad, CA, USA). The cells were stained for 15 min with an antibody against Ki-67 (clone B56, Alexa Fluor 647 conjugate, 5 μL) while the permeabilizer was present. The cells were then washed twice and resuspended in 0.125 mL of 2% paraformaldehyde for 20 min. After a final wash and resuspension with 125 μL PBS supplemented with 10% fetal bovine serum (fluorescence-activated cell sorting [FACS] buffer), all samples were run on a BD LSRII Flow Cytometer within 24 hrs. Flow data were analysed using Flowjo version 9.8.2. Absolute CD4+ T cell counts were determined by multiplying the absolute number of lymphocytes obtained by complete blood count by the percentage of lymphocytes that stained positive for CD3 and CD4 and negative for CD20 and CD8 by flow cytometry.

### Gut and lymph node histology

Tissues were collected from anesthetized macaques, fixed in 10% formalin, then embedded in paraffin (FFPE). Five micron thick sections were cut from FFPE blocks and mounted on charged slides. To remove paraffin, slides were baked at 80^o^C, treated with xylene (5 min x 3), and hydrated through graded alcohols to deionized water. Heat-induced epitope retrieval was performed in pH 9.0 Tris-EDTA solution (10mM tris base, 1 mM EDTA, 0.05% tween-20) for 3 minutes (Ki67) or 1 minute (CD4) in a Decloaking Chamber (Biocare Medical, Concord, CA). Slides were then rinsed with PBS and blocked with 0.3% H_2_O_2_ in PBS for 10 min at room temperature followed by serum [10% goat serum (Sigma, St. Louis, MO) in PBS] for 1 hr at room temperature. Slides were incubated with primary antibody in PBS with 1% goat serum and 0.1% Triton X-100 overnight at 4^o^C in dilutions of 1:800 for anti-Ki67 (BD Pharmingen, 556003) or 1:100 for anti-CD4 (Leica, NCL-L-CD4-368). Slides were then rinsed thrice with PBS and treated with Signal Stain Boost IHC Detection Reagent (HRP, Mouse) (Cell Signaling Technology, Beverly, MA) for 30 min at room temperature. Slides were then rinsed three times with PBS and treated with DAB substrate (Cell Signaling Technology, Beverly, MA) for 1 min and Mayer’s hematoxylin (Sigma, St. Louis, MO) for 1 min. Slides were then rinsed in H_2_O and dehydrated through graded alcohols to xylene.

Immunohistochemical (IHC) stains for Ki67 and CD4 were scored by a pathologist who was blinded to study design and treatment assignment. Quantification was reported as the average of 10 high-power fields (600X). In the lymph nodes, Ki67 positive cells were counted within the parafollicular region only, since germinal centers were universally positive. In the colon both Ki67 and CD4 were quantified within the lamina propria and crypt epithelial cells were excluded.

### Statistical analysis

Information on statistical tests used to determine significance can be found in corresponding figure legends. All statistical analyses were performed in Graphpad Prism software version 6.0h (GraphPad Software, Inc., La Jolla, CA).

## Acknowledgements

The authors thank the University of Wisconsin, Department of Pathology and Laboratory Medicine and the WNPRC for funding and the use of its facilities and services.

## Funding

This work was funded by NIH grant R01 AI116382-01. This publication was made possible in part by a grant (P51OD011106) from the Office of Research Infrastructure Programs (ORIP), a component of the National Institutes of Health (NIH), to the Wisconsin National Primate Research Center (WNPRC), University of Wisconsin-Madison. The authors thank Drs. Thomas Friedrich and Shelby O’Connor for use of their core services associated with WNPRC, as well as Dr. Eva Rakasz for her advice on immunophenotyping. This research was conducted in part at a facility constructed with support from Research Facilities Improvement Program grant numbers RR15459-01 and RR020141-01. IHC was performed by Joseph Hardin using a core facility affiliated with the University of Wisconsin Carbone Cancer Center Support Grant (P30 CA014520). ALB performed this work with support from the University of Wisconsin’s Medical Scientist Training Program (MSTP) (grant T32 GM008692). The content is solely the responsibility of the authors and does not necessarily represent the official views of the National Institutes of Health. The funders of this research had no role in study design, data collection and analysis, decision to publish, or preparation of the manuscript.

## Author contributions

ALB, CRB, and DHO conceived of the experimental design. CRB collected tissues and performed all flow cytometry experiments and analysis with assistance from ALB and MM. MB and LMS assisted with tissue collection. CRB performed the sequencing analysis. AJE helped prepare the SIVmac239 inoculum. EJP, KGB, and SCIII provided clinical support for infected animals, performed tissue biopsies, and administered virus inocula. HAS processed tissues for histopathological analysis. DM and DTY performed the histopathological analyses. ALB wrote the manuscript with edits, input, and approval from all authors.

## Competing interests

The authors declare no competing interests.

## Data and materials availability

The coding-complete genome sequence for the SPgV strain used in this study can be found in the Genbank database under accession number KF234530.

## Supplementary Materials

Supplemental figure: Fig S1

Supplemental Tables: Tables S1-S4

